# KDM5A enhances MHC-I antigen presentation and potentiates antitumor immunity unleashed by immune checkpoint blockade

**DOI:** 10.1101/2025.06.05.658174

**Authors:** Dan Li, Chi He, Zhaoli Tan, Tantan Wang, Hong Wu, Pengcheng Zuo, Dimiter S. Dimitrov, Liangliang Wang, Xuebin Liao

## Abstract

While some studies have implicated the histone 3 lysine 4 demethylase KDM5A in tumorigenesis and promotion of metastasis, KDM5A has also been shown to boost immunotherapy responses in melanoma. The distinct functional role of KDM5A in the context of immunotherapy and the underlying mechanisms remain largely unknown. Here, we report that higher KDM5A expression strongly correlates with higher MHC-I expression, CD8^+^ T cell infiltration, and prolonged survival in colorectal cancer and gastric cancer patients receiving anti– PD-1 therapy. KDM5A promotes MHC-I-mediated antigen presentation and CD8^+^ T cell-mediated tumor killing across multiple murine cancer models. Mechanistically, KDM5A upregulates MHC-I expression through the SOCS1/IFN-γ/STAT1 signaling and improves tumor cell-intrinsic antigen presentation capacity by inhibiting lysosomal proteases in a demethylase-dependent manner. Moreover, KDM5A directly represses lysosomal cathepsins in dendritic cells and promotes the cross-priming activity. Through a high-throughput chemical compound library-based screen, we identified Clomiphene, an FDA-approved small molecule, which significantly elevates KDM5A expression, and enhances antitumor immunity during anti–PD-1 immunotherapy in mouse models of melanoma and colon cancer. Our findings that KDM5A functions in MHC-I-mediated immune activation during anti–PD-1 therapy present an opportunity for developing KDM5A-enhancing therapies to increase tumor immunogenicity, sensitize solid tumors to immunotherapy and exclude immune evasion.

## INTRODUCTION

The histone 3 lysine 4 demethylase KDM5A (also referred to RBP2 or JARID1A) removes the di- and tri- methylation on lysine 4 of histone H3 (H3K4me2/3) near the transcription start site (TSS), leading to the represses of transcription for its target genes.^1^ Previously reported as an oncogene, KDM5A has been reported to transcriptionally modulate tumor suppressor genes associated with cancer cell proliferation, invasion, as well as the drug resistance in various cancers.^2,3^ Thus, several KDM5A inhibitors have been developed; however, these have only achieved limited effectiveness in preclinical studies.^4^ In seeming contrast, we previously demonstrated that KDM5A increases PD-L1 expression in tumor cells and improves the response to anti–PD-1 immunotherapy by remodeling the tumor immune microenvironment.^5^

Major histocompatibility complex class-I (MHC-I) molecules present antigens to CD8^+^ T cells and results in activation of antitumor immune responses.^6^ Dysregulation of MHC-I molecules has been linked to intrinsic and acquired resistance to immunotherapy in cancer patients.^7,8^ Genetic modifications (Ras/MAPK, NF-κB pathway)^9,10^ and epigenetic modulation (DNA methylation)^11^ have been shown to contribute to MHC-I expression in tumor cells. Recent studies have reported that epigenetic therapy can potentiate antitumor immunity through restoration of the MHC-I antigen processing and presentation machinery in both tumor cells and antigen presenting cells.^12–14^ However, the epigenetic regulation of MHC-I has not been explored in depth. Those findings have fueled our ongoing efforts to understand how KDM5A modulates MHC-I-mediated antitumor immunity during immunotherapy.

Here, immunohistochemical staining of pre anti–PD-1 immunotherapy biopsies revealed a positive correlation for KDM5A expression with MHC-I expression and with better ICB response in a patient cohort with colorectal cancer and gastric cancer. KDM5A increases IFN-γ-dependent MHC-I expression by directly repressing SOCS1 in a demethylase-dependent manner. KDM5A epigenetically reduces the transcription of lysosomal cathepsins, thereby promotes the antigen presentation activity of both tumor cells and DCs, respectively. Increasing KDM5A expression levels using an FDA-approved small molecule (Clomiphene) resulted in elevated CD8^+^ T cell-dependent antitumor immunity, and thus improved the responsivity to anti– PD-1 therapy in murine models of multiple cancers. Our study provides insights into how KDM5A regulates MHC-I expression and associated antigen presentation in both cancer cells and immune cells to sensitize tumors to ICB therapy.

## RESULTS

### Elevated KDM5A levels are associated with increased MHC-I and improved response responsivity to anti–PD-1 therapy in patients

To evaluate the potential relevance of KDM5A for clinical responses to anti–PD-1 therapy, we examined the KDM5A protein level through immunohistochemical staining of tumor tissues from colorectal and gastric cancer patients enrolled in a clinical trial at the Fifth Medical Center of 301 Hospital, Beijing, China (No. KY-2023-6-43); patients were treated with Sintilimab (anti–PD-1 antibody). Briefly, a high KDM5A level was strongly correlated with prolonged progression-free survival (PFS) in patients receiving the Sintilimab monotherapy (r=0.9224; **Figure 1A**) as well as in patients receiving the combination treatment (r=0.8509; **figure S1A**). Beyond examining clinical responses, we also conducted a Pearson correlation analysis and detected positive correlations between the KDM5A protein level and increased tumor-infiltration of CD8^+^ T cells (r=0.3267; **Figure 1B**) as well as the expression levels of multiple MHC-I molecules (r=0.6667; **Figure 1C-D**).

**Figure 1.**
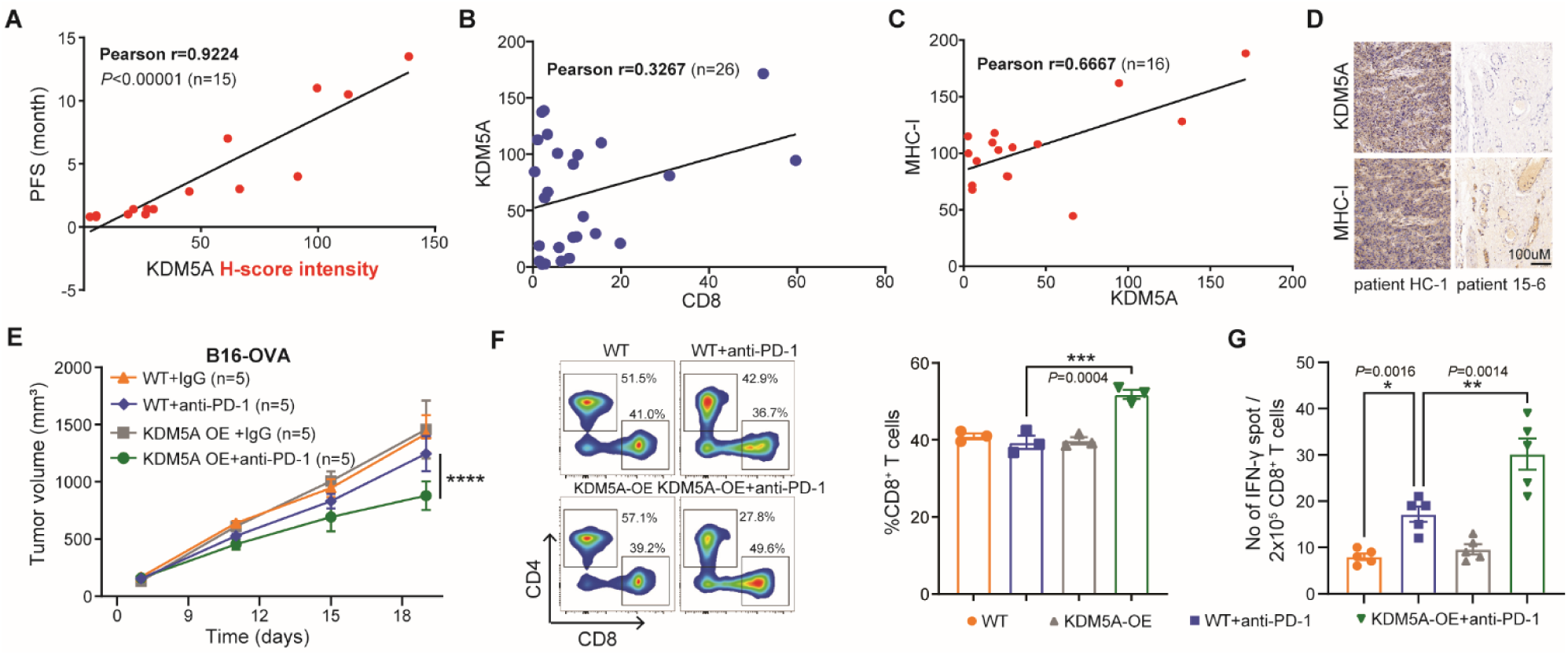
The H3K4 demethylase KDM5A positively correlates with better response to anti–PD-1 therapy in both humans and murine cancers. **(A)** Pearson correlation between KDM5A H-score intensity and the survival (PFS) of cancer patient receiving anti–PD-1 monotherapy (n=15). Biopsies were collected approximately two-four weeks after the anti–PD-1 therapy. **(B)** Pearson correlation between KDM5A H-score intensity and tumor infiltrating CD8^+^ T cell level of cancer patient receiving anti–PD-1 monotherapy (n=26). **(C)** Pearson correlation between KDM5A H-score intensity and tumor infiltrating CD8^+^ T cell level of cancer patient receiving anti–PD-1 monotherapy (n=16). **(D)** Tissue sections from patients immunohistochemically stained for KDM5A and MHC-I. Representative KDM5A-high (Patient HC-1) and KDM5A-low (Patient 15-6) specimens are shown. Scale bars, 100 μm. **(E)** WT mice were injected subcutaneously with 1×10^6^ WT or KDM5A-OE B16-OVA cells. When the tumor size reached 100 mm^3^, tumor-bearing mice were treated with anti–PD-1 antibody (2 doses per week, three doses total). Tumor growth was monitored (n=5 per group). **(F)** Representative flow cytometric analysis and quantification of CD8^+^ T cells in CD45^+^ tumor infiltrating lymphocytes (TILs) in B16-OVA tumors at day 18 of treatments (IgG control and anti–PD-1 antibody, n = 5 mice). **(G)** IFN-γ production in CD8^+^ T cells after coculturing with DCs. WT B16-OVA or KDM5A-OE B16-OVA tumors in WT mice were treated with anti–PD-1 antibody. Tumor draining lymph nodes were removed and digested to purify the CD11c^+^ DCs. The DCs were cocultured with CD8^+^ T cells from OT-I mice for 3 days and the cross-priming activity of DCs was analyzed by IFN-γ ELISPOT assay. Data are represented as mean ± s.e.m. One of two or three representative experiments was shown (E-G). Statistical analysis was performed using two-way ANOVA test with corrections for multiple variables (E), one-way ANOVA with Bonferroni’s multiple comparison tests (F, G). \**P* < 0.05, \*\**P* < 0.01, \*\*\**P* < 0.001, and \*\*\*\**P* < 0.0001.

In line with this observation, we used the Gene Expression Profile Interactive Analysis (GEPIA) database for public cancer data and found that *KDM5A* was positively correlated with multiple markers of CD8^+^ T-cell-dependent cytotoxic activity (*CD8A*, *GZMA*, *PRF1*, and *TNFA*)^15^ (**figure S1B**). Additionally, *KDM5A* was positively correlated with the extent of CD8^+^ T cell tumor infiltration in melanoma, prostate cancer, and kidney cancer patients (**figure S1C**). Analysis of transcriptomic datasets for cancer patients using the TIMER2.0 website^16^ revealed that *KDM5A* was positively correlated with an MHC-I gene signature (*B2M, HLA-A, HLA-B, HLA-C, HLA-E, HLA-F, TAP1*, and *TAP2*) in 35/40 examined cancer types (**figure S1D**).

MHC-I antigen processing machinery (APM) molecules have been reported to mediate responses to immunotherapy,^17,18^ we therefore investigated whether KDM5A is associated with MHC-I APM. *KDM5A* was correlated with the expression of many genes comprising a reported APM gene signature^19^ across the 40 examined tumor cohorts (**figure S1E**), including *ERAP1*, a gene encoding a protein known to function in MHC-I associated antigen processing and presentation^20, 21^. Notably, we found that *KDM5A* ranked first in the correlation of *ERAP1* with currently well-recognized histone modification regulators (**figure S1F**). These exploratory analyses consistently support impacts from KDM5A on antitumor immunity in cancer patients and implicate elevated expression of MHC-I and APM genes in these impacts.

### Overexpressing KDM5A enhances the antitumor response to anti**–**PD-1 therapy in murine cancers

To investigate whether the clinical impacts of elevated KDM5A observed in human cancers also occur in murine tumors models, we transplanted KDM5A-overexpresion (KDM5A-OE) B16-OVA cells into WT mice and treated the mice with the anti–PD-1 antibody. The anti–PD-1 therapy caused a more pronounced suppressive effect on the growth of KDM5A-OE tumors than on WT tumors (*P*<0.0001; **Figure 1E**), indicating that KDM5A in tumor cells sensitizes tumors to anti–PD-1 therapy in murine melanoma.

We subsequently investigated whether elevated KDM5A in tumors affects the cytotoxic T cell population by profiling the tumor-infiltrating immune cells using flow cytometry. Following anti–PD-1 treatment, the proportion of CD8^+^ T cells was significantly higher in KDM5A-OE tumors than in WT tumors (*P*=0.0004; **Figure 1F**). ELISPOT assays showed significantly increased IFN-**γ** production in anti–PD-1-treated CD8^+^ T cells sorted from KDM5A-OE tumors compared to those sorted from WT tumors (*P*=0.0014, **Figure 1G**). Further inspection of tumor infiltrating CD8^+^ T cells showed a decrease proportion of PD1^+^CD8^+^ T cells in “KDM5A-OE+anti–PD-1” tumors versus “WT+anti–PD-1” tumors (*P*=0.0034; **figure S2A**), suggesting less exhausted CD8^+^ T cells. No changes were observed in the numbers of the tumor infiltrating CD4^+^ T cells, NK cells, CD103^+^ DCs, or M1-type or M2-type macrophages (**figure S2B-D**). Thus, elevated KDM5A in tumors specifically induces more cytotoxic CD8^+^ T cells during anti–PD-1 therapy.

### KDM5A converts tumor cells into reprogrammed antigen presenting cells

Recalling the positive correlation detected between KDM5A and MHC-I in patient, we evaluated the expression of MHC-I and AMP genes in a set of *KDM5A* knockout (*KDM5A*-KO) and overexpression cancer cell lines. Briefly, compared to WT human breast MDA-MB-231 cells, knockout of *KDM5A* resulted in significantly reduced levels of MHC-I (total amount of HLA protein) (*P*<0.0001), while overexpression of KDM5A significantly increased MHC-I levels (*P*=0.0127) (**Figure 2A and figure S3A**); the same trends were observed in three murine cancer cell lines (4T-1, MC38, and B16-OVA) with similarly modulated *KDM5A* status (**Figure 2B and figure S3B**). A bulk RNA-seq analysis of the WT and *KDM5A*-KO MDA-MB-231 cells (and subsequent qPCR validation) showed that knocking out *KDM5A* resulted in significantly decreased expression of MHC-I associated genes including *HLA-A*, *HLA-B*, *HLA-C*, *B2M*, *TAP1*, and *TAP2* (**figure S3C-D**). Conversely, KDM5A-OE MDA-MB-231 and B16-OVA cells showed significantly increased expression of MHC-I and APM genes compared to the corresponding WT cells (**figure S3E**). These findings collectively support that KDM5A somehow regulates the expression of both MHC-I and APM genes in tumor cells.

**Figure 2.**
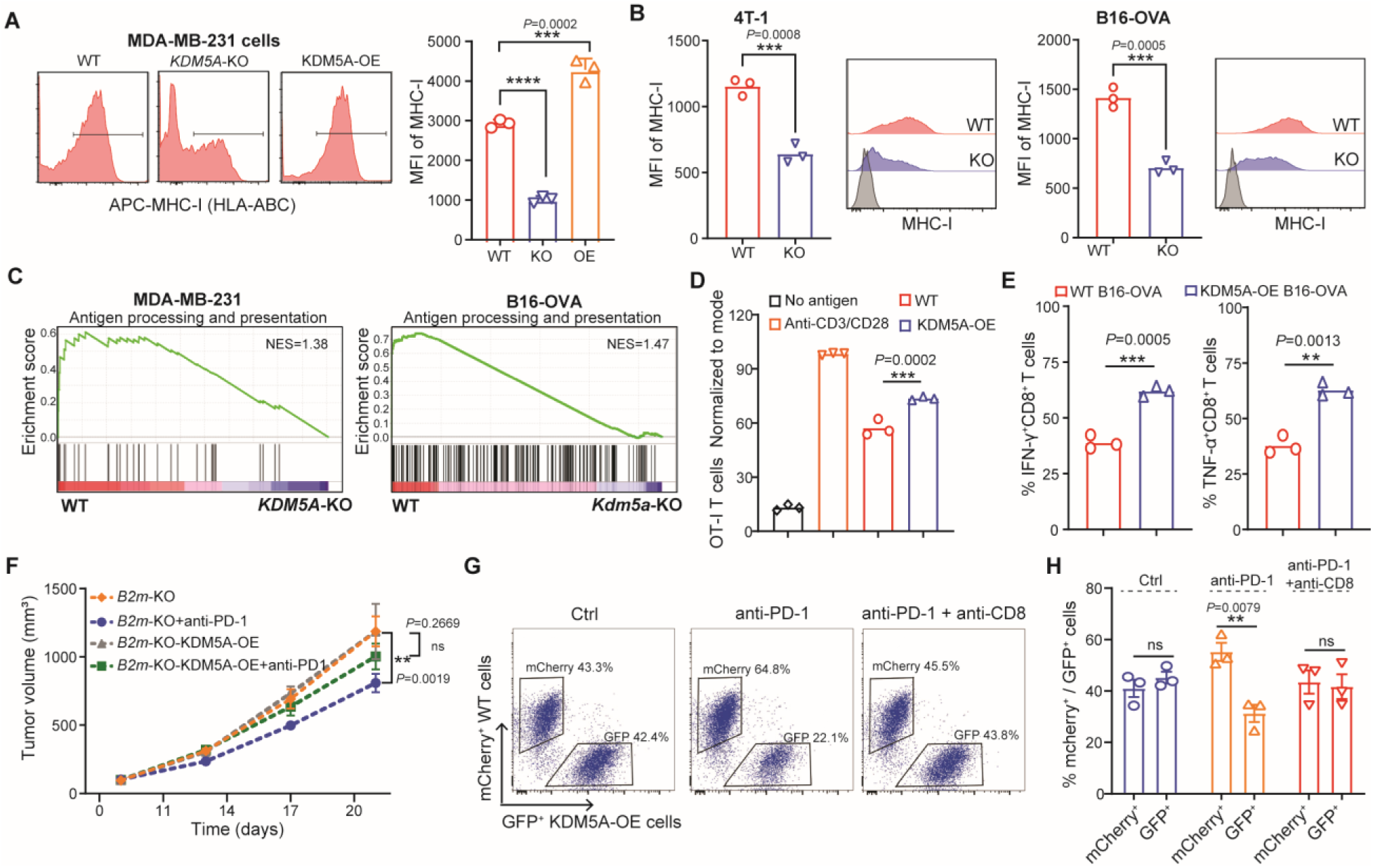
KDM5A increases the MHC-I expression and antigen presentation activity of tumor cells. **(A)** Representative flow cytometry analysis of MHC-I (left) and MFI (Mean fluorescent intensity) of MHC-I^+^ cells of WT, *KDM5A* knockout (KO) and KDM5A-overexpresion (OE) MDA-MB-231 cells (right). **(B)** Mean fluorescent intensity of MHC-I in WT, *Kdm5a* knockout (KO) 4T-1 and B16-OVA cells by flow cytometry. **(C)** GSEA plots of reactome antigen processing and presentation signaling showing positively correlation with KDM5A in WT (versus KDM5A knockout) MDA-MB-231 (left) and B16-OVA cells (right). NES, normalized enrichment score. **(D)** Flow cytometry analysis of an *in vitro* proliferation assay showing the frequency of proliferating OT-I CD8^+^ T cells when co-cultured with WT or KDM5A-OE B16-OVA cells. **(E)** Representative flow cytometry analysis of IFN-γ^+^ (or TNF-α^+^) OT-I CD8^+^ T cells in the co-culture system of WT or KDM5A-OE B16-OVA cells. **(F)** WT mice were injected subcutaneously with 1×10^6^ *B2m*-knockout or *B2m*-knockout KDM5A-OE B16-OVA cells. When the tumor size reached 100 mm^3^, tumor-bearing mice were treated with anti–PD-1 antibody (2 doses per week, three doses total). Tumor growth was monitored (n=5 per group) **(G-H)** Representative flow cytometry analysis of mcherry^+^ B16-OVA cells and GFP^+^ KDM5A-OE B16-OVA cells for ctrl, anti–PD-1 antibody treatment and anti–PD-1 antibody plus anti-CD8 antibody treatment (G). Ratio of mcherry^+^ B16-OVA cells: GFP^+^ KDM5A OE B16-OVA cells for conditions indicated (H). Data are represented as mean ± s.e.m. One of two or three representative experiments was shown (A, B, D-H). Statistical analysis was performed using one-way ANOVA with Bonferroni’s multiple comparison tests (A, D), two-sided unpaired Student’s *t*-test (B, E, H), or two-way ANOVA test with corrections for multiple variables (F). \**P* < 0.05, \*\**P* < 0.01, and \*\*\**P* < 0.001.

It seems plausible that the elevated MHC-I content in tumor cells may reflect enhanced antigen presentation activity, so we reanalyzed the RNA-seq data and found that the “antigen processing and presentation” pathway was enriched in both WT human MDA-MB231 and murine B16-OVA cells compared to the corresponding knockout cells (**Figure 2C**). We subsequently investigated the functional role(s) of KDM5A in tumor cell-mediated antigen presentation by co-culturing KDM5A-OE B16-OVA cells with naïve CD8^+^ T cells. Note that we used proliferation of antigen-specific CD8^+^ T cells as a readout of antigen presentation activity, as proliferating antigen-specific CD8^+^ T cells can be instructed to develop into cytotoxic T cells (CTLs)^22,23^. Overexpression of KDM5A led to an increased antigen presentation capability of B16-OVA cells, as evidenced by an improved CD8^+^ T cell proliferation (*P*=0.0002; **Figure 2D**). Consistently, the levels of both IFN-γ and tumor necrosis factor (TNF)-α in CD8^+^ T cells were markedly increased when co-cultured with KDM5A-OE cells (*P*=0.0005; **Figure 2E**), representing enhanced cytotoxic function. These findings indicate that KDM5A may directly convert cancer cells into reprogrammed antigen presenting cells, a scenario in line with the working principle of a recently reported strategy for immunotherapy^24^.

Finally, we used KDM5A-overexpressing (OE) *B2m* knockout (*B2m*-KO) B16-OVA cells, in which MHC-I protein expression was diminished, to perform syngeneic mouse model experiments with WT mice. The anti– PD-1 antibody had no effect on inhibiting *B2m*-KO-KDM5A-OE B16-OVA tumors (*B2m*-KO-KDM5A-OE *vs. B2m*-KO-KDM5A-OE+anti–PD-1, *P*=0.2669) (**Figure 2F**), observed in WT-KDM5A-OE tumors. In addition, we employed a competition assay using mcherry^+^ WT cells and GFP^+^ KDM5A-OE B16-OVA cells, and noticed that anti–PD-1 therapy preferentially induces KDM5A-OE cells killing (*P*=0.0079), and the ability was abolished by blocking CD8^+^ T cells (**Figure 2G-H**). Together, these results demonstrated that KDM5A enhances MHC-I-mediated antigen presentation activity and ensues tumor sensitivity to anti–PD-1 therapy *in vivo*.

### KDM5A epigenetically represses *SOCS1* and activates IFN-γ/STAT1/MHC-I signaling

We next sought to determine which downstream pathways of KDM5A are required for MHC-I expression in tumor cells, we reanalyzed our RNA-seq data for WT and *KDM5A*-KO MDA-MB-231 cells. Both GO and GSEA analysis indicated enrichment for the “JAK-STAT signaling pathway” in WT not in *KDM5A*-KO cells (**Figure 3A**). Considering that mouse and clinical studies have documented that IFN-γ can induce MHC-I expression in the tumor microenvironment,^25,26^ it is conceivable that KDM5A may regulate IFN-γ-induced MHC-I expression. We treated WT and *KDM5A*-KO cells with recombinant IFN-γ and measured MHC-I expression by flow cytometry. A lower level of MHC-I was observed in three *Kdm5a*-KO cells (MDA-MB-231, B16-OVA, and 4T-1 cells) compared to WT upon the stimulation of IFN-γ (**Figure 3B**). A qPCR analysis showed significantly decreased levels of nine MHC-I associated genes in both *KDM5A*-KO MDA-MB-231 and B16-OVA cells as compared to the corresponding WT control cells (**figure S4A**). These results demonstrated that KDM5A is required to IFN-γ-induced MHC-I expression.

**Figure 3.**
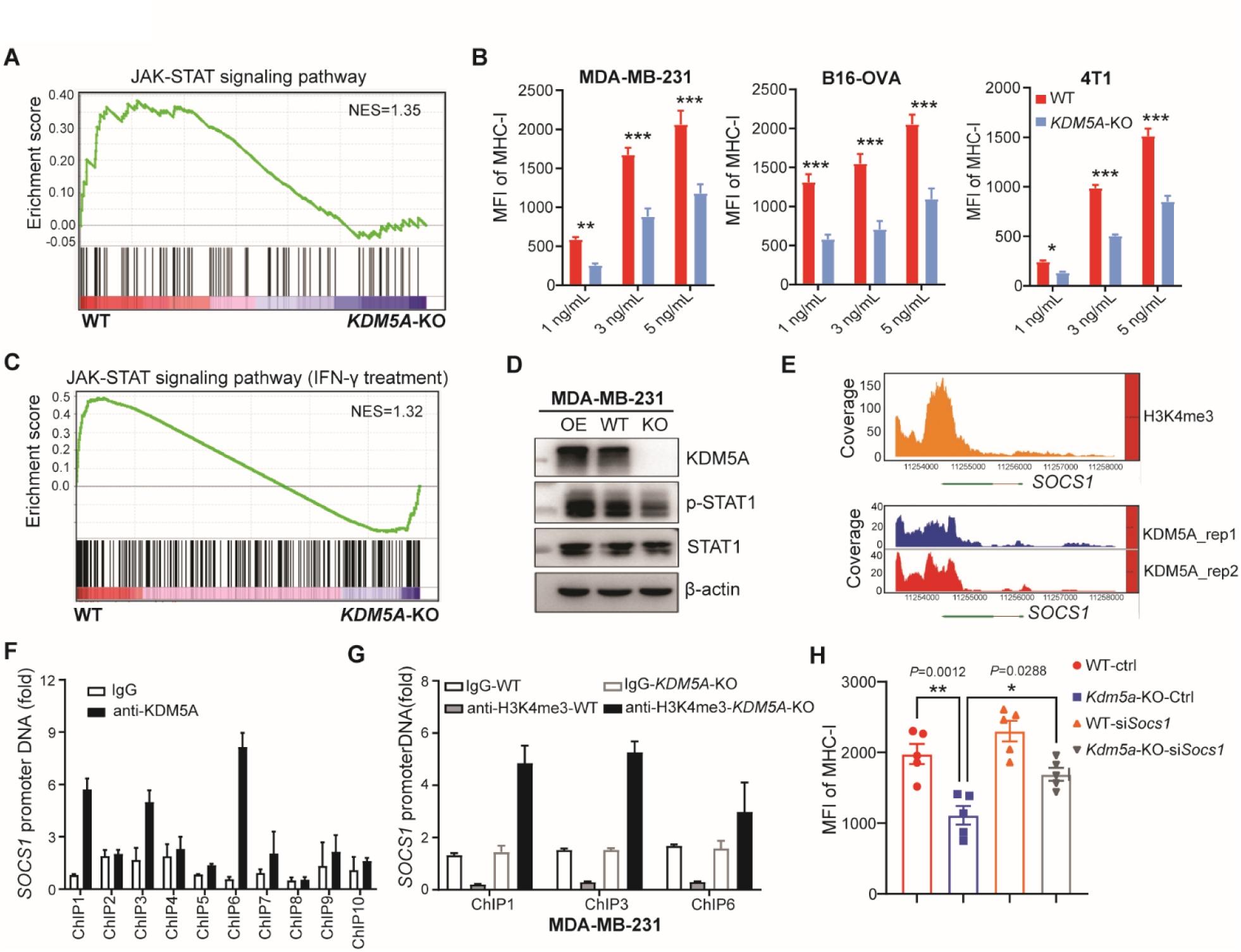
KDM5A binds at the *Socs1* promoter and decrease *Socs1* transcription to increase MHC-I expression in an IFN-γ-STAT1 dependent manner. **(A)** GSEA plots of JAK-STAT signaling showing positively correlation with KDM5A in WT MDA-MB-231 cells (versus *KDM5A*-KO cells). NES, normalized enrichment score. **(B)** Mean fluorescent intensity of MHC-I in WT and *KDM5A*-KO MDA-MB-231(HLA-ABC), 4T1 (H-2Kb) and B16-OVA (H-2Kb) cells treated with IFN-γ (1 ng/ml, 3 ng/ml and 5 ng/ml) for 24 hr by flow cytometry. **(C)** GSEA plots of JAK-STAT signaling showing positively correlation with KDM5A in IFN-γ (5 ng/ml for 24hr) treated WT MDA-MB-231 cells (versus *KDM5A*-KO cells). NES, normalized enrichment score. **(D)** Western blot analysis of STAT1 and p-STAT1 in IFN-γ (5 ng/ml for 24hr) treated WT, *KDM5A*-KO and KDM5A-OE MDA-MB-231 cells. **(E)** H3K4me3 binding peaks (top) and KDM5A binding peaks (bottom) across *Scos1* promoter in B16-OVA cells based on Cut & tag sequence **(F-G)** Chromatin immunoprecipitation (ChIP) analysis of the *SOCS1* promoter in MDA-MB-231 cells. KDM5A binding (**F**). H3K4me3 in WT and *KDM5A*-KO cells (**G**). **(H)** WT and *Kdm5a*-KO B16-OVA cells were transduced with siRNAs targeting Socs1. After 48-72 hr, the cells were subjected to flow cytometry analysis. Mean fluorescent intensity of MHC-I in these indicated four types of cells. Data are represented as mean ± s.e.m. One of two or three representative experiments was shown (A, B, D-H). Statistical analysis was performed using two-sided unpaired Student’s *t*-test (B), or one-way ANOVA with Bonferroni’s multiple comparison tests (H). \**P* < 0.05, \*\**P* < 0.01, and \*\*\**P* < 0.001.

We next performed RNA-seq of WT and *KDM5A*-KO MDA-MB-231 cells treated with IFN-γ. GSEA analysis revealed that KDM5A knockout mediated downregulation of genes sets related to “IFN-γ mediated signaling pathway” and “JAK-STAT signaling pathway” (**Figure 3C and figure S4B**). Such a scenario was consistently observed in an independent mouse cell line B16-OVA cells (*Kdm5a*-KO vs. WT) based on the

RNA-seq data (**figure S4C**). These data let us speculate that KDM5A may affect IFN-γ/JAK/STAT signaling in tumor cells. We thus conducted immunoblotting assay and confirmed that KDM5A-overexpression resulted in increased phospho-STAT1 levels in MDA-MB-231 cells (**Figure 3D**). KDM5A knockout decreased phospho-STAT1 in both cytoplasm and nuclear (**figure S4D**), demonstrating an impaired IFN-γ/STAT1 signaling. These results indicate that *Kdm5a* knockout represses MHC-I expression dependent of IFN-γ at both transcriptional and protein levels in human and murine tumor cells.

Given that KDM5A is a known H3K4me2/3 demethylase, we conducted CUT & tag sequencing of B16-OVA cells using a KDM5A specific antibody to map target loci bound by KDM5A. Integrative Genome Viewer indicated a good fit between the H3K4me3 peaks and the KDM5A-binding peaks in the promoter region of *Socs1* (**Figure 3E**), a well-documented negative regulator of p-STAT1 activation^27^. ChIP-qPCR analysis supported that KDM5A directly binds to the *Socs1* promoter region (∼3.0 kb proximal to the transcription start site) (**Figure 3F**). Of note, ChIP-qPCR analysis showed the amount of elevated H3K4me3 levels at the *Socs1* promoter in the *KDM5A*-KO cells compared to WT cells (**Figure 3G**). Moreover, the *Socs1* mRNA level was significantly increased in *KDM5A*-KO cells compared to WT cells (**figure S4E**). Our data indicate that KDM5A reduces the transcription of *Socs1* in an H3K3me3-depednent manner.

We next sought to determine whether the regulation of MHC-I expression by KDM5A requires SOCS1. To achieve this, we generated *Socs1* knockdown cells using the corresponding siRNA in WT and *Kdm5a*-KO B16-OVA cells, and conducted flow cytometry analysis of MHC-I. The downregulation of MHC-I was observed in *Kdm5a*-KO cells compared to WT cells (*P*=0.0012), while the level of MHC-I was drastically increased when *Kdm5a*-KO cells were treated with *Socs1*-siRNA (*P*=0.0288) (**Figure 3H**). Collectively, these findings indicate that KDM5A epigenetically reduces SOCS1, and thereby activates IFN-γ/JAK/STAT1/MHC-I signaling.

### KDM5A represses the transcription of lysosomal cathepsins and increases the antigen presentation activity of tumor cells

To investigate the mechanisms by which KDM5A promotes antigen presentation activity of tumor cells, we performed Cistrome-GO analysis with our CUT & Tag-seq for B16-OVA cells. KDM5A-targeted genes were enriched for four lysosome-associated pathways (**Figure 4A**). Limiting lysosomal proteolysis can enhance antigen degradation and presentation^28,29^. We noticed that obvious peaks were present in the promoter regions of *Ctsb*, *Ctsf*, and *Ctsl*, signifying the direct binding (**Figure 4B**). The immunoblotting assay showed that *Kdm5a* knockout increased CTSB, CTSF, and CTSZ abundance in MDA-MB-231 cells, increased CTSL in B16-OVA cells (**Figure 4C**), while KDM5A overexpression decreased the expression of these CTS proteins (**Figure 4D**). These data demonstrate that KDM5A directly targets and down-regulates a group of lysosomal cathepsins-encoding genes in tumor cells.

**Figure 4.**
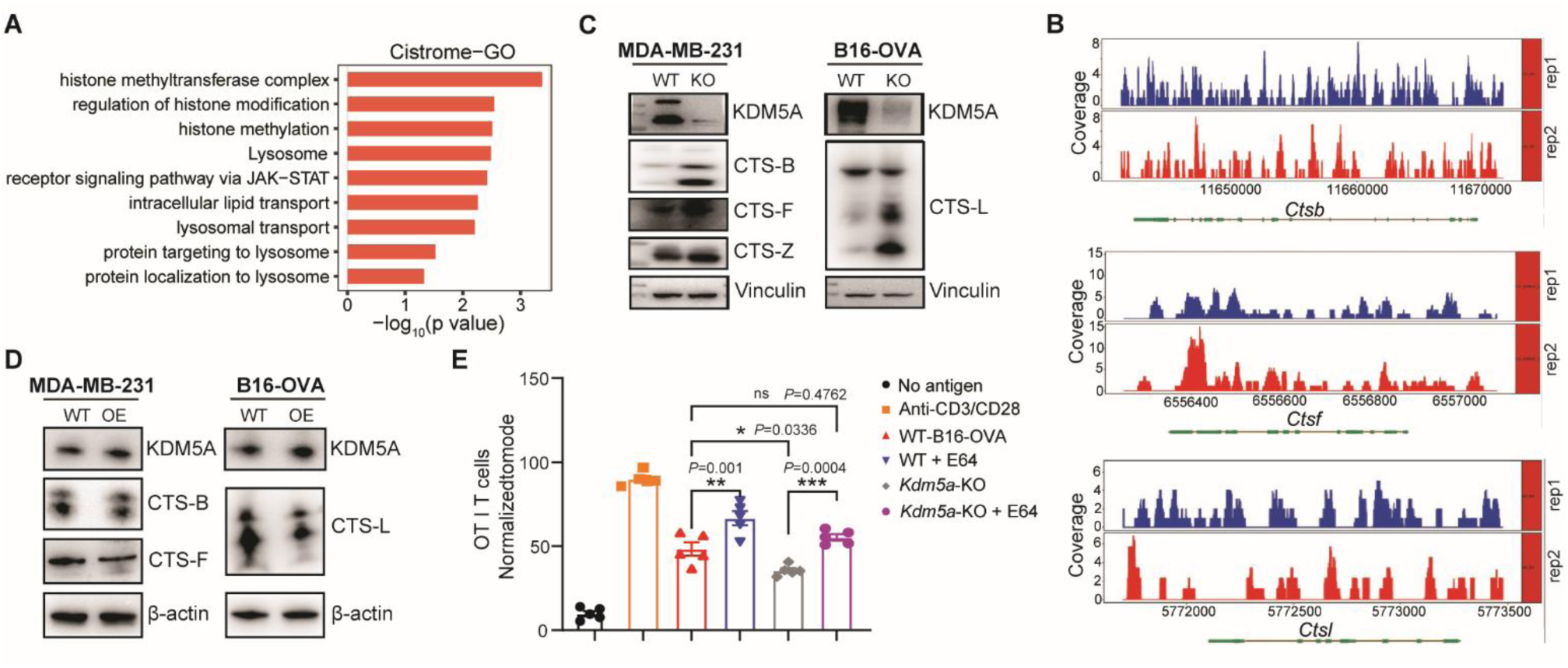
KDM5A represses the lysosomal cathepsins to increase the antigen presentation activity of tumor cells. **(A)** Function enrichment analysis (Cistrome-GO) of enriched genes in B16-OVA cells based on CUT &Tag seq data. **(B)** KDM5A binding ability at the promoter of *Ctsb*, *Ctsf*, and *Ctsl* in WT B16-OVA cells based on the CUT & Tag analysis. **(C)** Western blot analysis of CTSB, CTSF, CTSZ, and CTSL in WT and *KDM5A* knockout MDA-MB-23 (left) and *Kdm5a*-KO B16-OVA cells (right). **(D)** Western blot analysis of CTSB, CTSF and CTSL in WT and KDM5A-OE MDA-MB-231 cells (left) and B16-OVA cells (right). **(E)** WT or *Kdm5a*-KO B16-OVA cells were pre-treated with 2 μM E64 and co-cultured with OT-I CD8^+^ T cells. Flow cytometry analysis of proliferating OT-I CD8^+^ T. Data are represented as mean ± s.e.m. One of two or three representative experiments was shown (C-E). Statistical analysis was performed using one-way ANOVA with Bonferroni’s multiple comparison tests (E). \**P* < 0.05, \*\**P* < 0.01, and \*\*\**P* < 0.001.

We next to further demonstrate that the effect of KDM5A on antigen presentation capacity of tumor cells is dependent on lysosomal proteases. We treated B16-OVA cells with broad-spectrum cysteine protease inhibitor E64 and then co-cultured them with naïve CD8^+^ T cells. The inhibitor E64 significantly enhanced the efficiency of antigen presentation of WT B16-OVA cells (*P*=0.001; **Figure 4E**). Notably, although *Kdm5a*-KO B16-OVA cells exhibited attenuated antigen presentation function compared to WT B16-OVA cells (*P*=0.0336), inhibitor E64 can rescue this to the baseline level (compared to WT B16-OVA, *P*=0.4762, **Figure 4E**). Taken together, these data highlight that KDM5A reduces the transcription of lysosomal cathepsins in tumor cells and promotes tumor cell-intrinsic antigen presentation capacity.

### KDM5A reduces the transcription of lysosomal cathepsins and induces the cross-priming capacity of DCs

Given our finding in patients for correlation of KDM5A and MHC-I levels, and considering the strong KDM5A expression reported for tumor infiltrating DCs in melanoma patients treated with ICB therapy^30^ (**figure S5A**), we speculate that KDM5A may drive MHC-I expression in DCs as it does in tumor cells. To test this, we isolated DCs from CD11c^Cre+^*Kdm5a*^fl/fl^ conditional knockout mice (hereafter *Kdm5a*-cKO) and measured MHC-I expression by flow cytometry. *Kdm5a* knockout had no effects on MHC-I expression in both cDC1 and cDC2 cells (**figure S5B**).

We subsequently assessed whether KDM5A contributes the MHC-I mediated cross-priming activity by co-culturing isolated DCs with activated naïve CD8^+^ T cells. *Kdm5a*-cKO DCs exhibited apparently reduced tumor associated antigen presentation efficiency relative to WT DCs, evidenced by a significantly lower level of IFN-γ (*P*<0.0001; **Figure 5A**). Moreover, *Kdm5a*-cKO DCs cross-presented less tumor-associated antigen to OT-I CD8^+^ T cells compared to WT-DCs, in terms of lower proliferation rate (**Figure 5B and S5C-D**). We next excluded the role of KDM5A on DC activation markers since *Kdm5a* knockout did not affect LPS, PolyI:C, or R848-induced expression of MHC-II, CD80, CD86 and IL12p40 (**figure S5E-G**). Collectively, our findings indicate that loss of *Kdm5a* decreases the cross-priming capacity of DCs.

**Figure 5.**
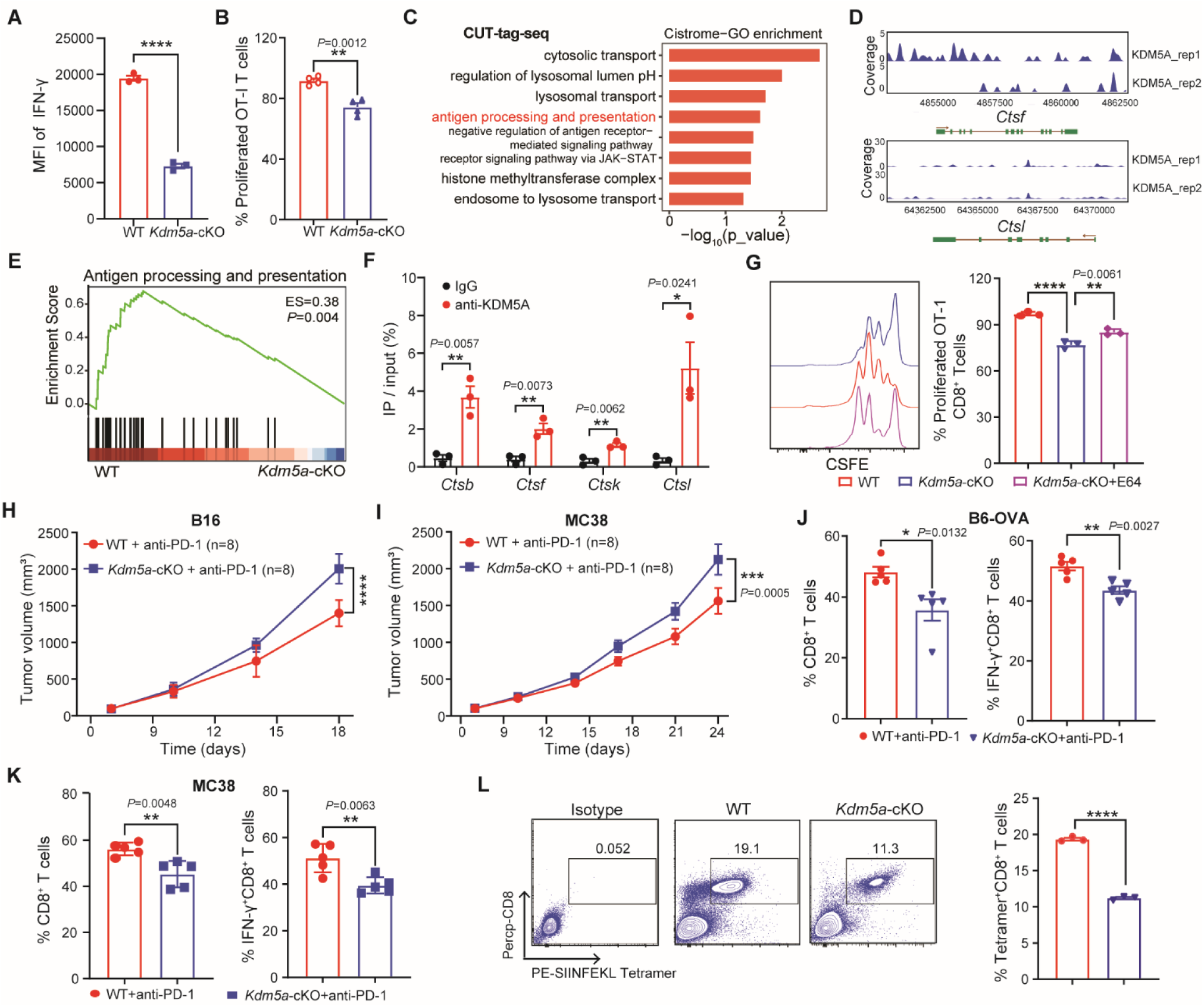
*Kdm5a* deficiency in DCs attenuates cross-priming capacity and reduces the response to anti–PD-1 therapy. **(A)** WT or *Kdm5a*-cKO CD11c^+^DCs were isolated from TDLN in B16-OVA tumor bearing mice (day 19) and were subjected to co-cultured with naïve CD8^+^ T cells for 72h. Intracellular IFN-γ^+^ staining in CD8^+^ T cells were analyzed by flow cytometry. **(B)** OT-I CD8^+^ T cell proliferation stimulated by WT cDC1s and *Kdm5a*-cKO cDC1s with tumor associated antigen (TAA). **(C)** Function enrichment analysis (Cistrome-GO) of enriched genes in BMDC cells based on CUT & Tag seq data. **(D)** KDM5A binding ability at the promoter of *Ctsf*, and *Ctsl* in WT BMDCs based on the CUT & Tag analysis. **(E)** GSEA plots of reactome “Antigen processing and presentation” showing positively correlation with KDM5A in WT versus *Kdm5a*-cKO BMDCs based on RNA-seq data. **(F)** Chromatin immunoprecipitation (ChIP) analysis of the *Ctsb*, *Ctsf*, *Ctsk*, and *Ctsl* promoters in BMDCs with anti-KDM5A antibody. **(G)** *In vitro* OT-I CD8^+^ T cell proliferation stimulated by WT, *Kdm5a*-cKO and *Kdm5a*-cKO plus E64 BMDCs with soluble OVA antigen **(H-I)** Wild-type (*Kdm5a*^fl/fl^) or *Kdm5a*-cKO (*CD11c*^cre^ *Kdm5a*^fl/fl^) mice were injected subcutaneously with 1×10^6^ B16-OVA cells (H) or MC38 cells (I). When the tumor size reached 100 mm^3^, tumor-bearing mice were treated with anti–PD-1 antibody (2 doses per week, three doses total). Tumor growth was monitored (n=8 mice per group) **(J-K)** Flow cytometry analysis and quantification of total CD8^+^ T cells and IFN-γ^+^CD8^+^ T cells in CD45^+^ TILs in B16-OVA (J) and MC38 (K) tumors at day 18 of treatments. **(L)** Representative flow cytometry analysis of SIINFEKL tetramer^+^CD8^+^ T cells in the CD45^+^ tumor infiltrating lymphocytes (TILs) in B16-OVA tumors at day 18 of indicated treatment. Data are represented as mean ± s.e.m. One of two or three representative experiments was shown. Statistical analysis was performed using two-sided unpaired Student’s *t*-test (A, B, F, J, K, L), one-way ANOVA with Bonferroni’s multiple comparison tests (G), or two-way ANOVA test with corrections for multiple variables (H, I). \**P* < 0.05, \*\**P* < 0.01, \*\*\**P* < 0.001, and \*\*\*\**P* < 0.0001.

To investigate the potential downstream pathways of KDM5A induction of cross-priming capacity in DCs, we conducted CUT & tag sequencing of DCs to map target genes bound by KDM5A. The Cistrome-GO enrichment analysis showed that KDM5A-targeted genes were enriched for pathways associated with lysosomes and antigen processing and presentation (**Figure 5C**). Particularly, we found that the KDM5A-binding peaks were evident in the promoter region of *Ctsf*, and *Ctsl* (**Figure 5D**). Moreover, we performed RNA-seq of WT and *Kdm5a*-cKO DCs and the consequent GSEA and GO analysis. The “MHC-I antigen processing and presentation”-associated pathways were enriched in WT DCs (**Figure 5E and figure S6A**). qPCR analysis showed that a series of genes contributing to the antigen processing and presentation were down-regulated upon *Kdm5a* knockout in DCs (**figure S6B**). These data suggests that KDM5A promotes antigen processing and presentation activity of DCs likely due to the impaired lysosomal cathepsins.

To further demonstrate the relationship between KDM5A and lysosomal cathepsins-encoding genes, we perform ChIP-qPCR analysis with bone marrow derived DCs (BMDCs). We found that KDM5A directly binds to the promoter region of *Ctsb*, *Ctsf*, *Ctsk*, and *Ctsl* (**Figure 5F**), accompanied by a consistent decrease in transcription (**figure S6C**). At protein levels, *Kdm5a* knockout resulted in the up-regulated CTSF and CTSL in DCs (**figure S6D-E**). To strengthen the cause-effect relationship between KDM5A-mediated CTS proteins and cross-priming capacity of DCs, we treated *Kdm5a-*cKO BMDCs with protease inhibitor E64 and co-cultured with CD8^+^ T cells. Compared with WT BMDCs, *Kdm5a*-cKO BMDCs resulted in impaired antigen presentation function (*P*<0.0001), whereas inhibitor E64 can rescue such decrease (*Kdm5a*-cKO**+**E64 *vs. Kdm5a*-cKO, *P*=0.0061; **Figure 5G**). These data indicated that KDM5A enhances the cross-priming capacity of DCs by directly repressing lysosomal cathepsins.

### *Kdm5a* deficiency in DCs attenuates the response to anti**–**PD-1 therapy in murine cancers

To investigate whether *Kdm5a* deficiency in DCs could alter the response to ICB, we treated tumor-bearing *Kdm5a*-cKO mice and *Kdm5a*^fl/fl^ (hereafter WT) mice with anti–PD-1 antibody. We found that in the syngeneic murine melanoma (B16-OVA) model, anti–PD-1 treatment resulted in a diminished inhibition of tumor growth in *Kdm5a*-cKO mice compared with WT mice (*P*<0.0001; **Figure 5H**). Consistent tumor inhibition phenotype was observed in MC38 murine colon cancer model (*P*=0.0005; **Figure 5I**).

We subsequently depicted the shifts in the tumor microenvironment in *Kdm5a*-cKO mice by profiling the tumor-infiltrating immune cells with flow cytometry. Immune infiltrates contained lower levels of total CD8^+^ T cells and cytotoxic CD8^+^ T cells in tumors from *Kdm5a*-cKO mice than from WT mice following anti–PD-1 treatment (**Figure 5J**), suggesting the reduced immunosurveillance ability. Such a scenario is consistent with our results in MC38 tumors (**Figure 5K**). Accordingly, there was no obvious difference in the number of Tregs, DCs, NKs and B cells (**figure S6F**). To determine whether antigen-specific CD8^+^ T cell responses are involved, we analyzed the frequency of tumor-infiltrating CD8^+^ T cells expressing the SIINFEKL MHC-I tetramer and found a substantial decrease, almost by half, in *Kdm5a*-cKO tumors (*P*<0.0001; **Figure 5L**), indicating the impaired CD8^+^ T cell priming function. These findings demonstrate that loss of *Kdm5a* in DCs impairs the response to anti–PD-1 therapy dependently of CD8^+^ T cells.

### Pharmacological increase of KDM5A by Clomiphene improves the response to anti**–**PD-1 therapy

To demonstrate that enhanced KDM5A can be translated into a clinically relevant strategy, we screened an in-house compound library with by high-throughput RNA-seq–based high-throughput screening (HST2) combined flow analysis^31^. We identified that clomiphene, an FDA approved drug to treat female infertility with its no reported toxic effects^32^, increased the abundance of both KDM5A and MHC-I in tumor cells (**figure S7A-E**). Clomiphene also increased KDM5A expression in DCs in a dose dependent manner *in vitro* (**figure S7F**). We next examined whether clomiphene could improve the anti–PD-1 responses in murine cancer models. B16-OVA tumor-bearing mice were intratumorally treated daily with clomiphene starting on the day of anti–PD-1 treatment. Clomiphene treatment alone did not inhibit tumor growth, however, when combined with anti–PD-1 treatment, it significantly enhanced tumor growth inhibition compared with anti–PD-1 treatment alone (*P*<0.0001; **Figure 6A**). In terms of systemically delivery of clomiphene (i.g. injection), the combination treatment with clomiphene and anti–PD-1 also resulted in a pronounced inhibition of B16-OVA tumor growth, exerting an improved effect of ICB (**Figure 6B**). Our results demonstrate that combination therapy of clomiphene and anti–PD-1 rise to the robust antitumor effects.

**Figure 6.**
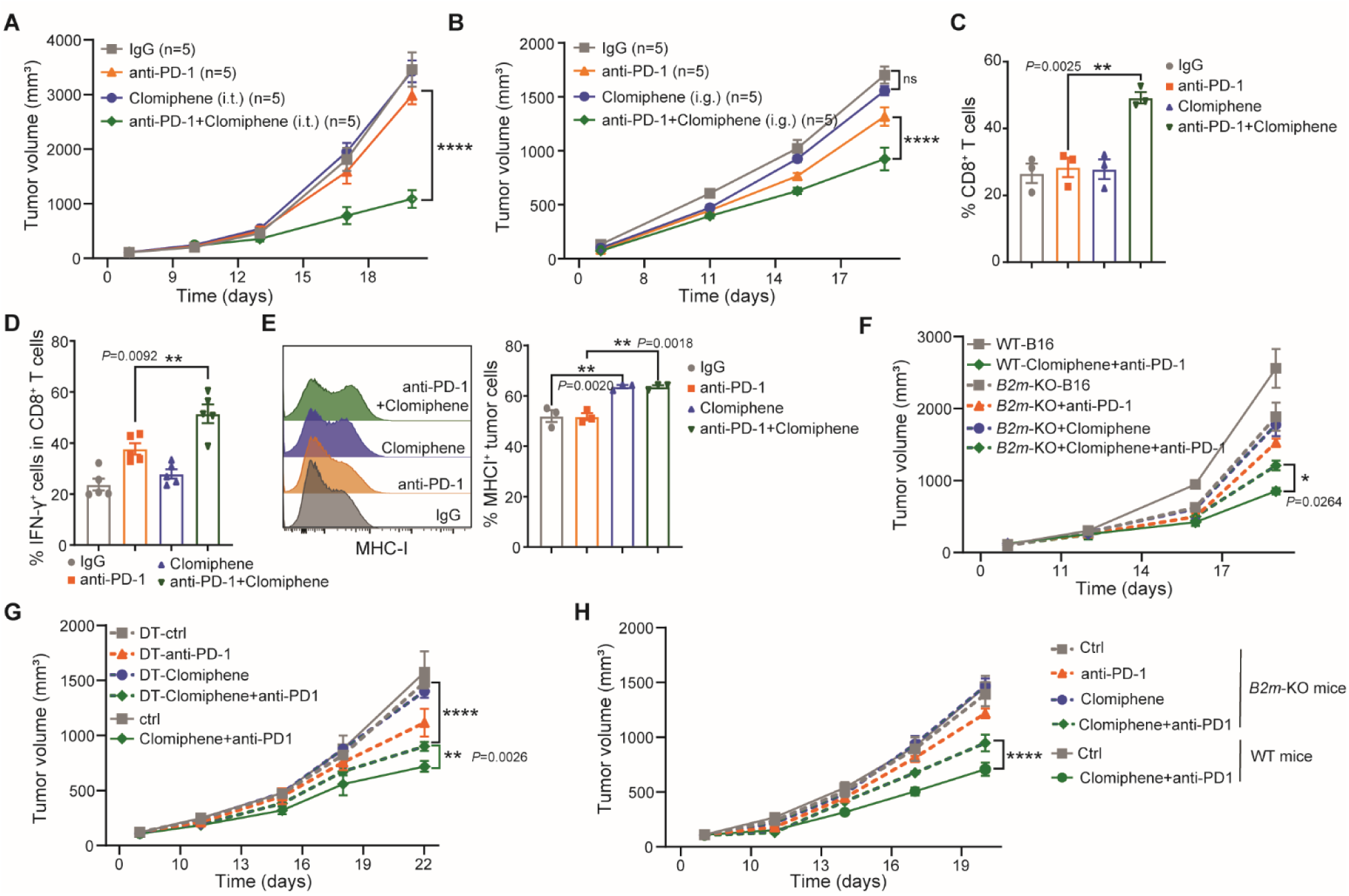
Clomiphene increases KDM5A expression and improves the efficacy of anti–PD-1 immunotherapy in melanoma murine model. **(A)** WT mice were injected subcutaneously with 1×10^6^ B16-OVA cells. When the tumor size reached 100 mm^3^, tumor-bearing mice were treated with anti–PD-1 antibody (2 doses per week, three doses total), and/or introtumorally (i.t.) injected with clomiphene. Tumor growth was monitored (n=5 per group). **(B)** B16-OVA tumor bearing mice were treated with anti–PD-1 antibody (2 doses per week, three doses total), and/or clomiphene (i.g. injection; 100 mg/kg). Tumor growth was monitored (n=5 per group) **(C-D)** Flow cytometry analysis and quantification of total CD8^+^ T cells (C) and IFN-γ^+^CD8^+^ T cells (D) in B16-OVA tumors with indicated treatments. **(E)** Flow cytometry analysis of MHC-I on CD45^-^ cells from B16-OVA tumors with indicated treatments. **(F)** WT B16-OVA or *B2m*-KO B16-OVA tumor bearing mice were treated with anti–PD-1 antibody (2 doses per week, three doses total), and/or clomiphene (i.g. injection; 100 mg/kg). Tumor growth was monitored (n=5 per group). **(G)** CD11c-DTR bone marrow chimeric mice were injected subcutaneously with 1 x10^6^ B16-OVA cells. When the tumor size reached 100 mm^3^, mice were injected with Diphtheria toxin (50 ng/ per mice). Next day, the mice were treated with anti–PD-1 antibody (2 doses per week, three doses total), and/or clomiphene. Tumor growth was monitored (n=5 per group). **(H)** WT mice or *B2m*-KO mice were injected subcutaneously with 1×10^6^ B16-OVA cells. When the tumor size reached 100 mm^3^, tumor-bearing mice were treated with anti–PD-1 antibody (2 doses per week, three doses total), and/or introtumorally (i.g. injection; 100 mg/kg) injected with clomiphene. Tumor growth was monitored (n=5 per group). Data are represented as mean ± s.e.m. One of two or three representative experiments was shown. Statistical analysis was performed using two-way ANOVA test with corrections for multiple variables (A, B, F, G, H), or one-way ANOVA with Bonferroni’s multiple comparison tests (C, D, E). \**P* < 0.05, \*\**P* < 0.01, \*\*\**P* < 0.001, and \*\*\*\**P* < 0.0001.

To explore the underlying immunological mechanisms, we profiled the tumor-infiltrating immune cells in B16-OVA tumors following the combination treatment with anti–PD-1 treatment (i.p.) and clomiphene (i.g.). We observed that the combination treatment significantly increased the number of tumor-infiltrating CD8^+^ T cells and CTL compared to anti–PD-1 treatment alone (*P*=0.0025, *P*=0.0092 respectively; **Figure 6C-D**). The numbers of tumor-infiltrating PD-1^+^CD8^+^ T cells were markedly decreased upon combination treatment (**figure S7G**). There was no significant difference in tumor-infiltrating Tregs, NK, CD103^+^ DCs, M1-type and M2-type macrophages (**figure S7H**). In according with our *in vitro* phenotype, clomiphene significantly increased MHC-I abundance on tumor cells (CD45^-^ cells) in B16-OVA tumors (**Figure 6E**) and increased the antigen-presentation ability of tumor cells (**figure S7I**). In pursuing this further, we inoculated *B2m*-KO B16-OVA cells into WT mice and treated the mice with clomiphene together with anti–PD-1 antibody. The antitumor efficacy of the combination treatment was abolished in *B2m*-KO tumors (**Figure 6F**). These data indicate that clomiphene improves the response to anti–PD-1 treatment relying on the increased CD8^+^ T cells and tumor cell-intrinsic MHC-I.

To investigate whether DCs are essential in the increased tumor growth control of the combination treatment with clomiphene and anti–PD-1, we employed the bone marrow chimeric mice with bone marrow cells (BMCs) from CD11c-DTR mice, whose DCs were depleted after diphtheria toxin (DT) treatment, for above-mentioned tumor growth experiments. Upon administration of DT, the combination treatment showed weaker tumor controlling ability compared to those without DT (*P*=0.0026; **Figure 6G**), suggesting the requirement of DCs. Such a similar impaired tumor growth inhibition phenotype was observed in *B2m* knockout mice (*P*<0.0001; **Figure 6H**), in which the MHC-I associated antigen presentation was destroyed. Collectively, these results demonstrate that the antitumor effect of the combination treatment with clomiphene and anti–PD-1 therapy depends on elevated KDM5A and MHC-I associated antigen presentation activity in both tumor cells and DCs, and is similar to that observed in KDM5A-overexpressing tumors. As an FDA approved anti-tumor drug, clomiphene demonstrates high translational potential in expanding therapeutic approaches to improve clinical outcomes of anti-cancer therapies. Taking into regard the aforementioned positive correlation between KDM5A and better clinical outcome in patients receiving Sintilimab monotherapy, our findings strongly imply the urgent clinical practices combing clomiphene and anti-PD-1 immunotherapy (as a promising strategy) against colorectal and gastric cancers, in which unmet medical needs are in high demands.

## DISCUSSION

Here we report that enhancing KDM5A improves immune checkpoint blockade efficacy by up-regulating MHC-I expression in tumor cells as well as promoting antigen presentation activates of both tumor cells and dendritic cells. KDM5A converts cancer cells into reprogrammed antigen presenting cells via two different mechanisms including: directly inhibits SOCS1 expression and promotes IFN-γ/STAT1 activation for MHC-I regulation; directly suppresses lysosomal cathepsins to increase antigen presentation activity. In DCs, KDM5A also targets and inhibits the transcription of cathepsins encoding genes and results in enhanced cross-priming capacity. Our study delineates a deeper link between KDM5A and MHC-I antigen presentation during immunotherapy.

It has been reported that the lower levels of KDM5A in human tumors is associated with immune exclusion and resistance to immune checkpoint blockade.^4^ We speculate that the deceased KDM5A in human tumors might be another MHC-I down-regulation/dysregulation mechanism, helping to explain that some patients do not benefit from ICB therapy. We show the evidence that KDM5A epigenetically increases tumor-intrinsic MHC-I expression in murine melanoma, colon cancer, breast cancer and human breast cancer. This is opposite to that KDM5A down-regulates the antigen presentation pathway in ovarian cancer.^33^ The regulation mechanism of MHC-I by KDM5A may be cancer type-specific. Further investigations are needed to clarify the regulation of MHC-I in relation to KDM5A in different cancer types.

Previous studies proposal that KDM5A promotes the progression of multiple solid tumors including lung cancer, breast cancer, and prostate cancer etc,^4,34,35^ contradicts the fact that KDM5A improves the response to ICB therapy in melanoma^5^. We further illustrate that enhancing KDM5A can improve the efficacy of anti–PD-1 immunotherapy in murine cancer models of melanoma and colon cancer; KDM5A positively correlated with better anti–PD-1 response in colorectal cancer and gastric cancer patients. These suggest that KDM5A may play a unique role in the response to anti–PD-1 therapy induced stress. Our results provide proof-of-principle preclinical evidence that enhancing KDM5A with clomiphene in a whole-animal setting notably improves the antitumor efficacy of anti–PD-1 immunotherapy. Further studies are needed to evaluate the pro and con effects of using this FDA-approved drug as a combination agent with anti–PD-1 immunotherapy for solid tumors treatment in the future.

## Acknowledgments

This work was funded by the National Natural Science Foundation of China (NSFC) (no. 21877067), Tsinghua-Peking Centre for Life Science, and Roche Postdoc Fellowship Program. We thank Prof. Jun Liu at Peking University for helpful discussion.

## Author Contributions

X.L. conceived and supervised research. D.L., C.H., and L.W. performed most of the experiments. Z.T., provided clinical trial samples. T.W. performed the bioinformatics analysis. H.W. and P.Z. helped to perform experiments. L.W., and D.S.D. contributed to extensive discussions. X.L. and D.L. designed the study. L.W., D.L., and X.L. wrote the manuscript. L.W. and X.L. edited the manuscript.

---

**Figure S1.**
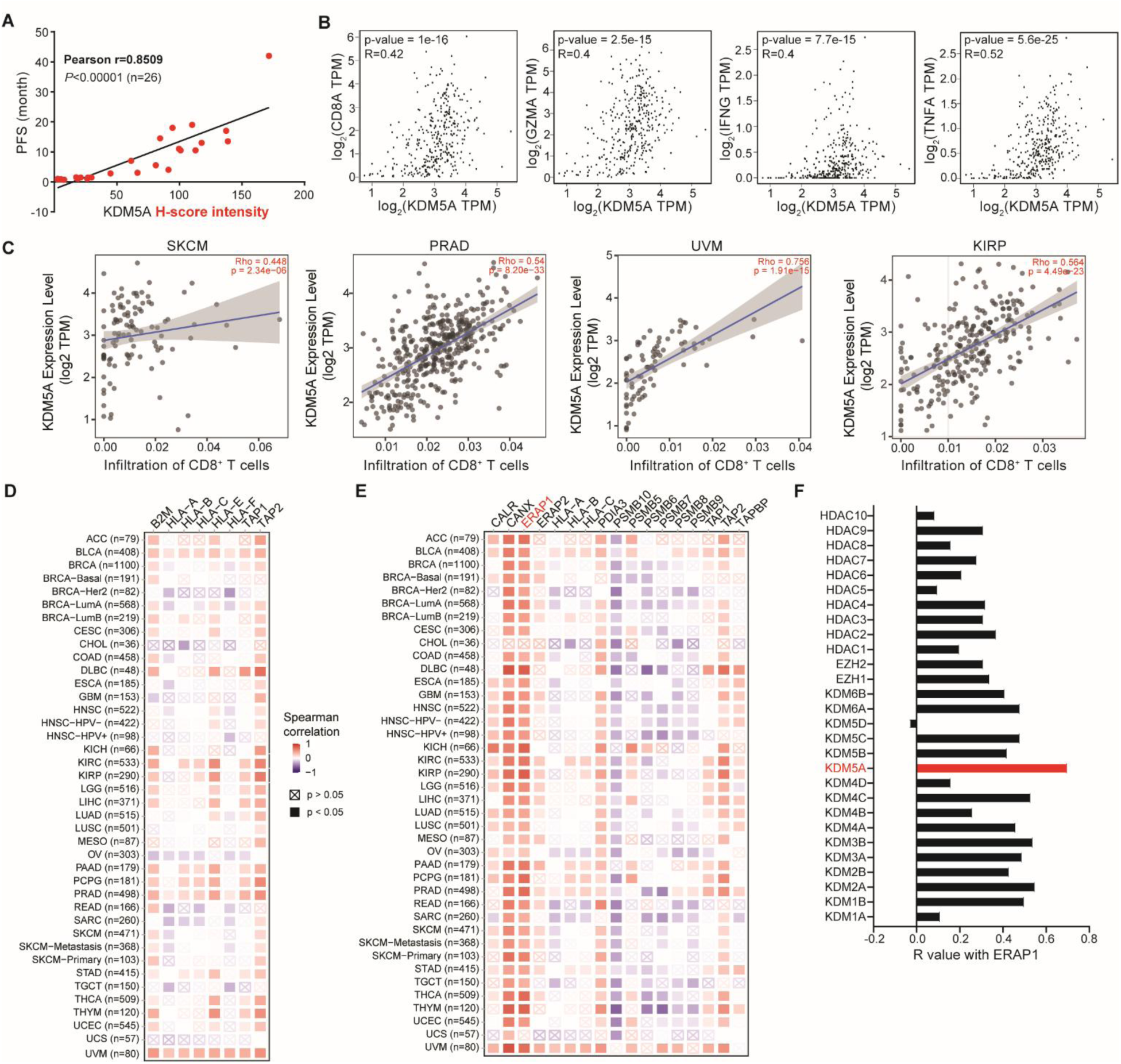
KDM5A positively correlates with immune response and MHC-I expression in TCGA tumor cohorts. Related to Figure 1. **(A)** Pearson correlation between KDM5A H-score intensity and the survival (PFS) of cancer patient receiving anti-PD-1 or combined with other therapy. **(B)** Scatter plots for the expression of KDM5A and CD8A, IFNG, GZMA and TNFA in melanoma and colon cancer (n=354). **(C)** Correlation of BAMBI expression and infiltration level of CD8^+^ T cells in skin cutaneous melanoma (SKCM), Prostate adenocarcinoma (PRAD), Uveal Melanoma (UVM), and Kidney renal clear cell carcinoma (KIRC) cohorts, obtained by TIMER2.0. R represents Spearman’s coefficient. **(D)** Heatmap showing the correlation between KDM5A and each gene in an MHC-I gene signature (*B2M*, *HLA-A*, *HLA-B*, *HLA-C*, *HLA-E*, *HLA-F*, *TAP1* and *TAP2*), obtained by TIMER2.0. (Spearman’s coefficient). **(E)** Heatmap showing the correlation between KDM5A and each gene in an MHC-I antigen processing machinery (APM) gene signature including 17 genes as indicated, obtained by TIMER2.0. (Spearman’s coefficient). **(F)** Spearman rank correlation coefficients between the ERAP1 expression and histone modification regulators as indicated.

**Figure S2.**
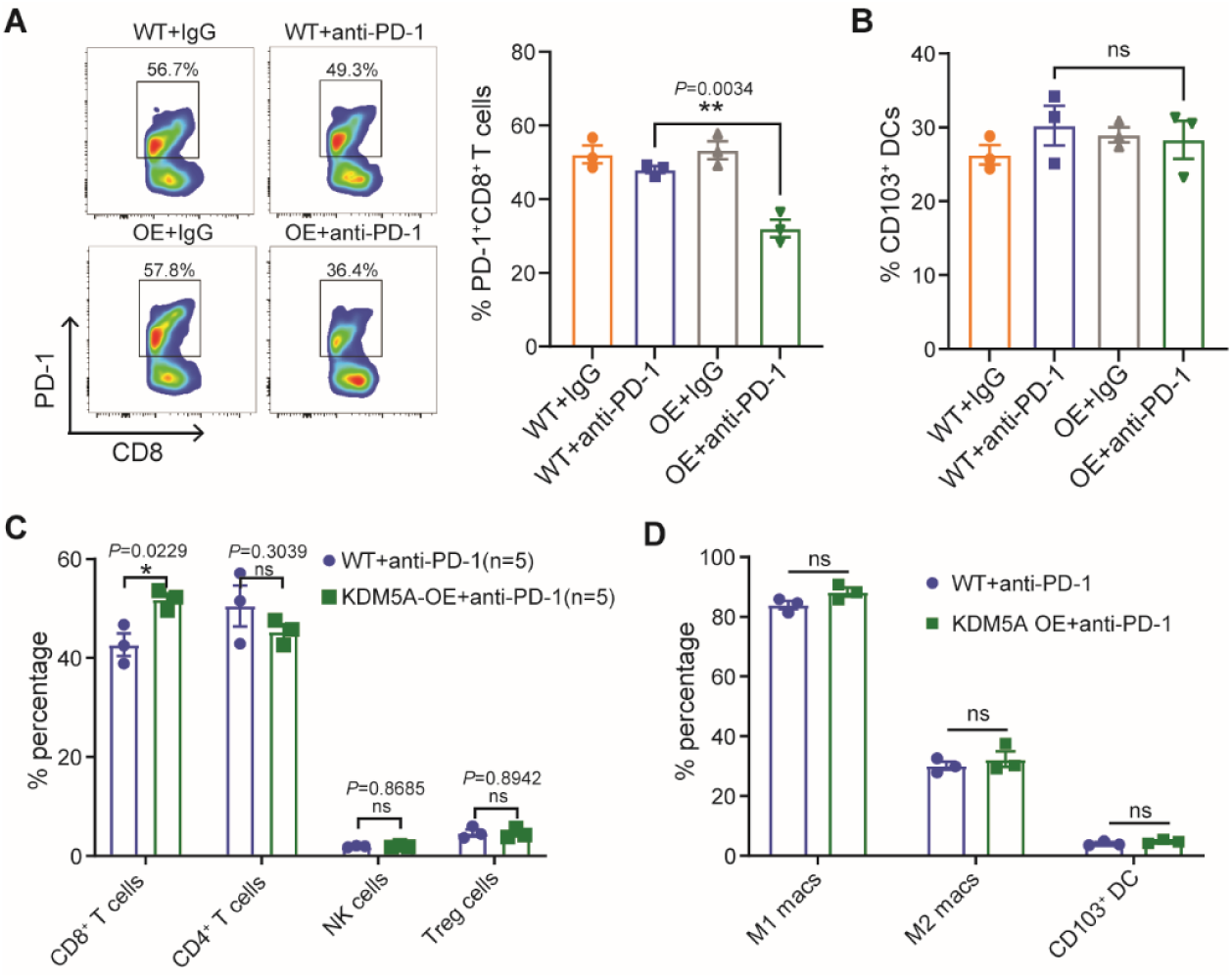
KDM5A overexpression has no significant effects on tumor infiltration of CD4^+^ T cells, NK cells, Macrophages and DCs in anti-PD-1 treated murine melanoma. Related to Figure 1. **(A)** Representative flow cytometric analysis and quantification of PD-1^+^CD8^+^ T cells in B16-OVA tumors at day 18 after anti-PD-1 treatments. **(B)** Representative flow cytometric analysis and quantification of CD103^+^ dendritic cells (DCs) in TME of B16-OVA tumors at day 18 after anti-PD-1 treatments. **(C)** Quantification by flow cytometry analysis of CD8^+^ T cells, CD4^+^ T cells, NK cells and Treg cells abundance in TME from B16-OVA tumor-bearing mice receiving the indicated treatments. **(D)** Quantification by flow cytometry analysis of M1-like macrophage, M2-like macrophage, and CD103^+^ DCs abundance in TME from B16-OVA tumor-bearing mice receiving the indicated treatments. Data are represented as mean ± s.e.m. One of two or three representative experiments was shown (A-D). Statistical analysis was performed using one-way ANOVA with Bonferroni’s multiple comparison tests (A, B) or two-sided unpaired Student’s *t*-test (C, D). \**P* < 0.05, \*\**P* < 0.01, \*\*\**P* < 0.001, and \*\*\*\**P* < 0.0001.

**Figure S3.**
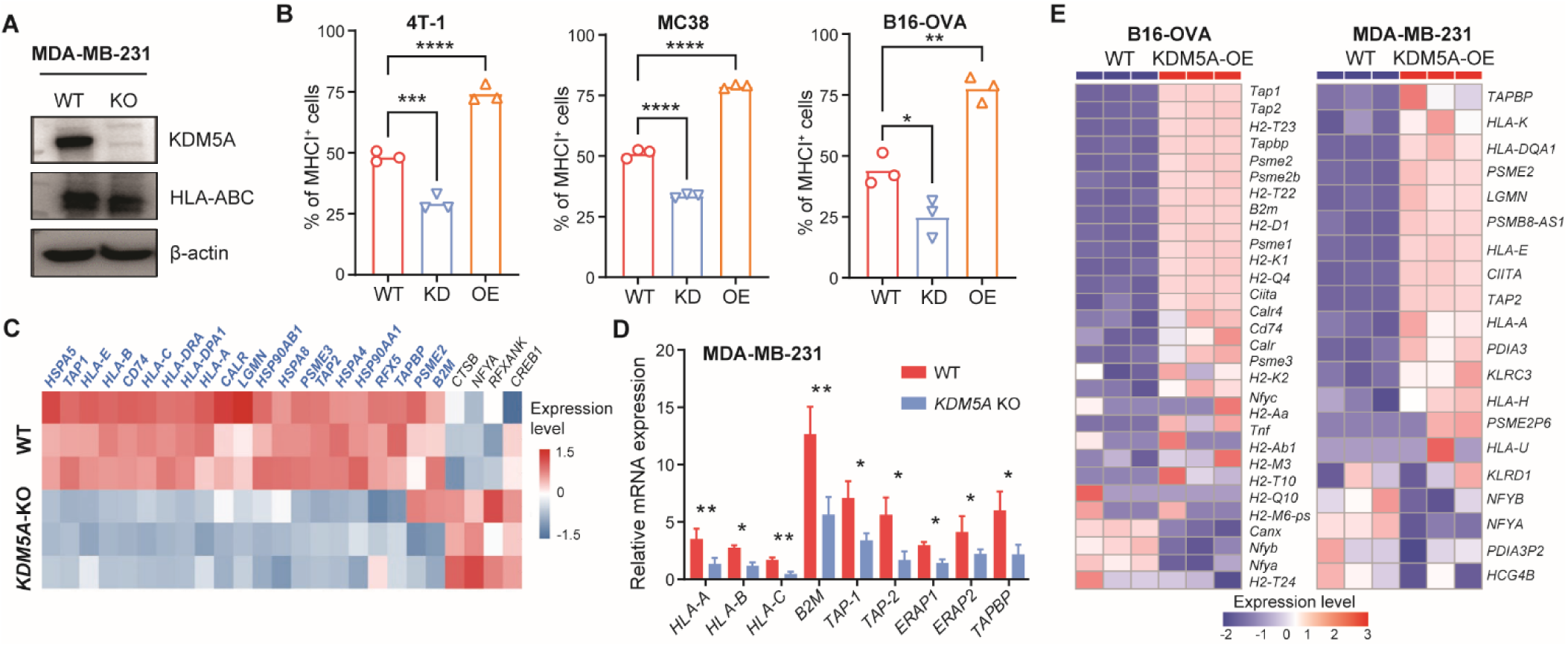
KDM5A increases the expression of MHC-I and APM genes in human and murine cancer cell lines. Related to Figure 2. (A) Western blot analysis of HLA-ABC in WT and *KDM5A*-KO MDA-MB-231 cells. **(B)** Representative flow cytometric analysis and quantification of MHC I (H-2Kb) in WT, *Kdm5a*-KD and KDM5A OE 4T-1, MC38 and B16-OVA cells. **(C)** Heatmap of a hierarchical clustering of the mRNA level of genes involved in the “antigen processing and presentation” signaling pathway. **(D)** Quantitative PCR (q-PCR) analysis of multiple genes associated with the MHC-I-APM in WT and *KDM5A*-KO MDA-MB-231 cells. **(E)** Heatmap of a hierarchical clustering of the mRNA level of genes involved in the “antigen processing and presentation” pathway in the indicated B16-OVA and MDA-MB-231 cells (WT *vs.* KDM5A-OE cells). Cluster analysis was performed with DAVID bioinformatics resources Data are represented as mean ± s.e.m. One of two or three representative experiments was shown (A, B, D). Statistical analysis was performed using one-way ANOVA with Bonferroni’s multiple comparison tests (B), or two-sided unpaired Student’s *t*-test (D). \**P* < 0.05, \*\**P* < 0.01, \*\*\**P* < 0.001, and \*\*\*\**P* < 0.0001.

**Figure S4.**
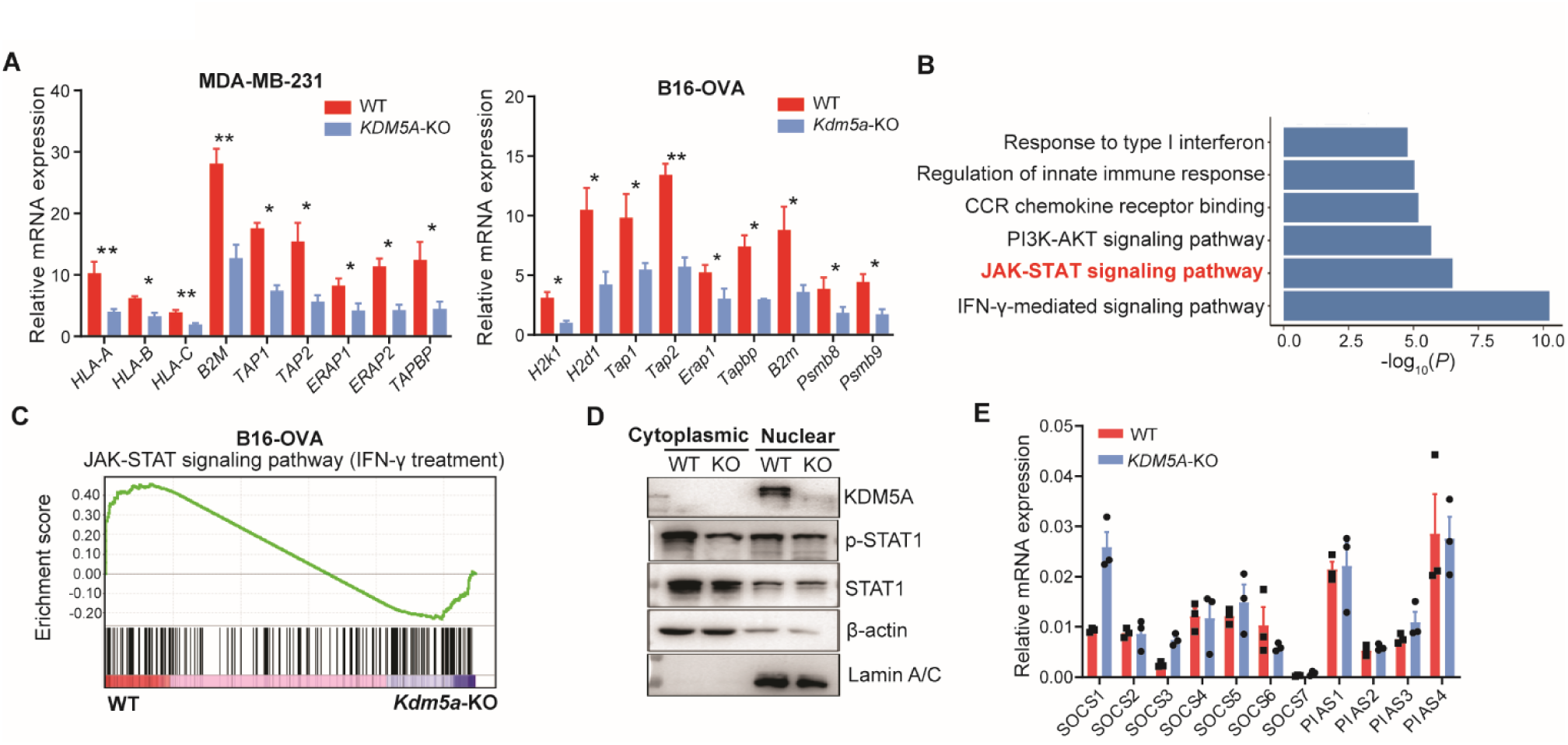
KDM5A inhibits *Socs1* transcription to activates IFN-γ/STAT1 signaling required for MHC-I expression in tumor cells. Related to Figure 3. **(A)** Quantitative PCR (q-PCR) analysis of multiple components of the MHC-I APM-associated genes in IFN-γ treated WT and *KDM5A*-KO MDA-MB-231 cells (left) or B16-OVA cells (right). **(B)** KEGG pathway analysis of WT and *KDM5A*-KO MDA-MB-231 cells based on the RNA-seq data. **(C)** GSEA plots of JAK-STAT signaling showing positively correlation with higher expression of KDM5A in IFN-γ treated B16-OVA cells. NES, normalized enrichment score. **(D)** Western blot analysis of STAT1 and p-STAT1 in cytoplasmic and nuclear of WT and *Kdm5a*-KO B16-OVA cells treated with IFN-γ. **(E)** qPCR analysis of mRNA expression for relative negative regulation genes involved in JAK-STAT pathway in *KDM5A*-KO compared with WT MDA-MB-231 cells. The qPCR data were normalized to *β-actin*. Data are represented as mean ± s.e.m. One of two or three representative experiments was shown (A, D, E). Statistical analysis was performed using two-sided unpaired Student’s *t*-test (A, E). \**P* < 0.05, and \*\**P* < 0.01.

**Figure S5.**
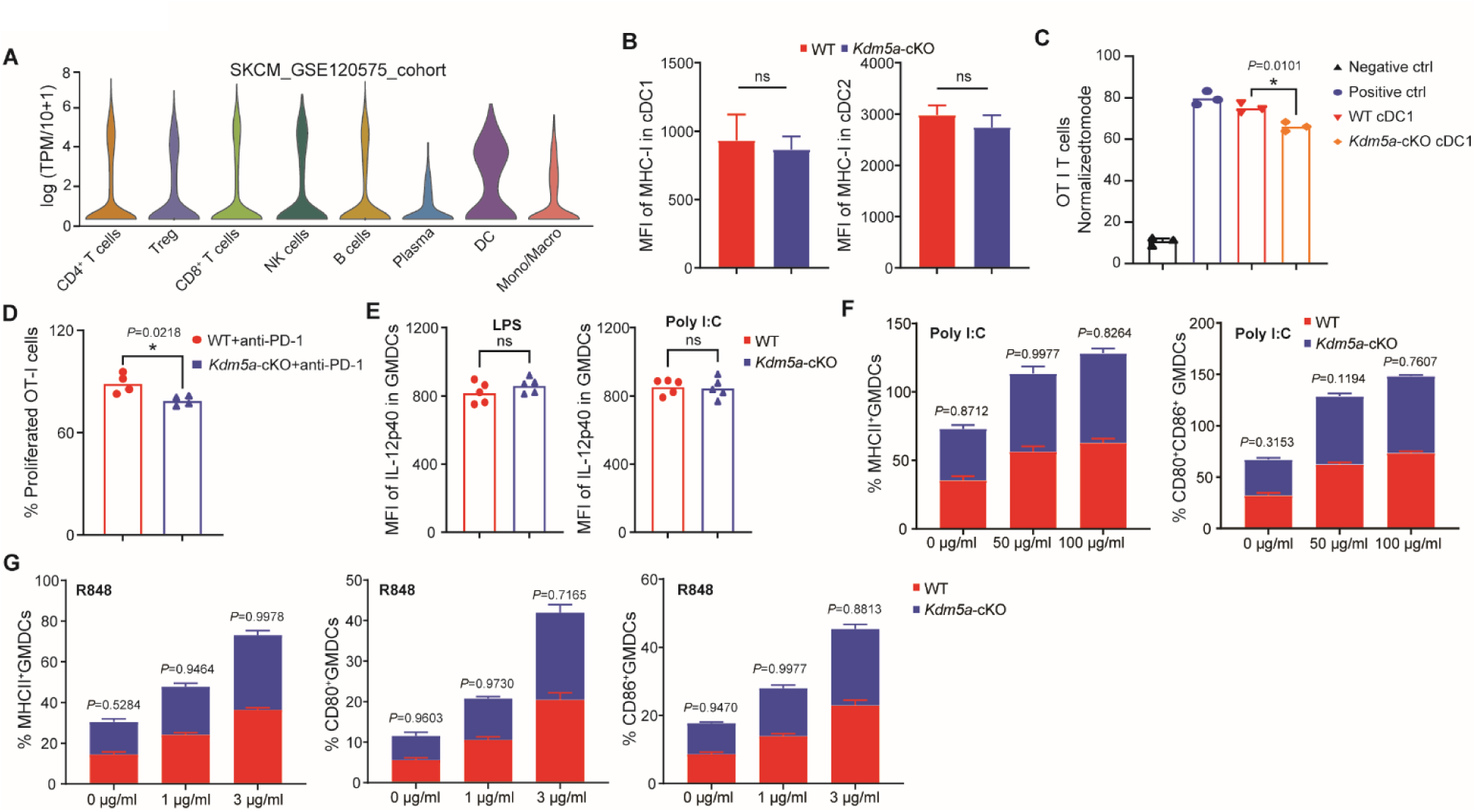
*Kdm5a* deficiency did affect the expression of MHC-I and activation markers in DCs. Related to Figure 5. **(A)** Violin plot showing the mRNA levels of KDM5A in different immune cells based on the public scRNA-seq data of human skin cutaneous melanoma (SKCM) (GSE120575), obtained by TISCH2. **(B)** Quantification by flow cytometry analysis of MHC-I in cDC1 and cDC2 from WT and *Kdm5a*-cKO mice. **(C)** *In vitro* OT-I CD8^+^ T cell proliferation stimulated by no antigen (negative control), anti-CD3/anti-CD28 (positive control), WT cDC1s and *Kdm5a*-cKO cDC1s with soluble antigen. **(D)** *Ex vivo* OT-I CD8^+^ T cell proliferation stimulated by WT cDCl and *Kdm5a*-cKO cDC1 with tumor associated antigen (TAA). **(E)** Representative flow cytometry analysis and quantification of IL-12p40 in LPS (100 ng/mL) and Poly I: C (50 ug/mL) treated BMDCs derived from WT and *Kdm5a*-cKO mice. **(F)** Quantification by flow cytometry analysis of MHC II, CD80 and CD86 in Poly I:C treated BMDCs from WT and *Kdm5a*-cKO mice. **(G)** Quantification by flow cytometry analysis of MHC II, CD80 and CD86 in R848 treated BMDCs from WT and *Kdm5a*-cKO mice Data are represented as mean ± s.e.m. One of two or three representative experiments was shown. Statistical analysis was performed using two-sided unpaired Student’s *t*-test (B, D-G) or one-way ANOVA with Bonferroni’s multiple comparison tests (C). \**P* < 0.05, \*\**P* < 0.01, and \*\*\**P* < 0.001.

**Figure S6.**
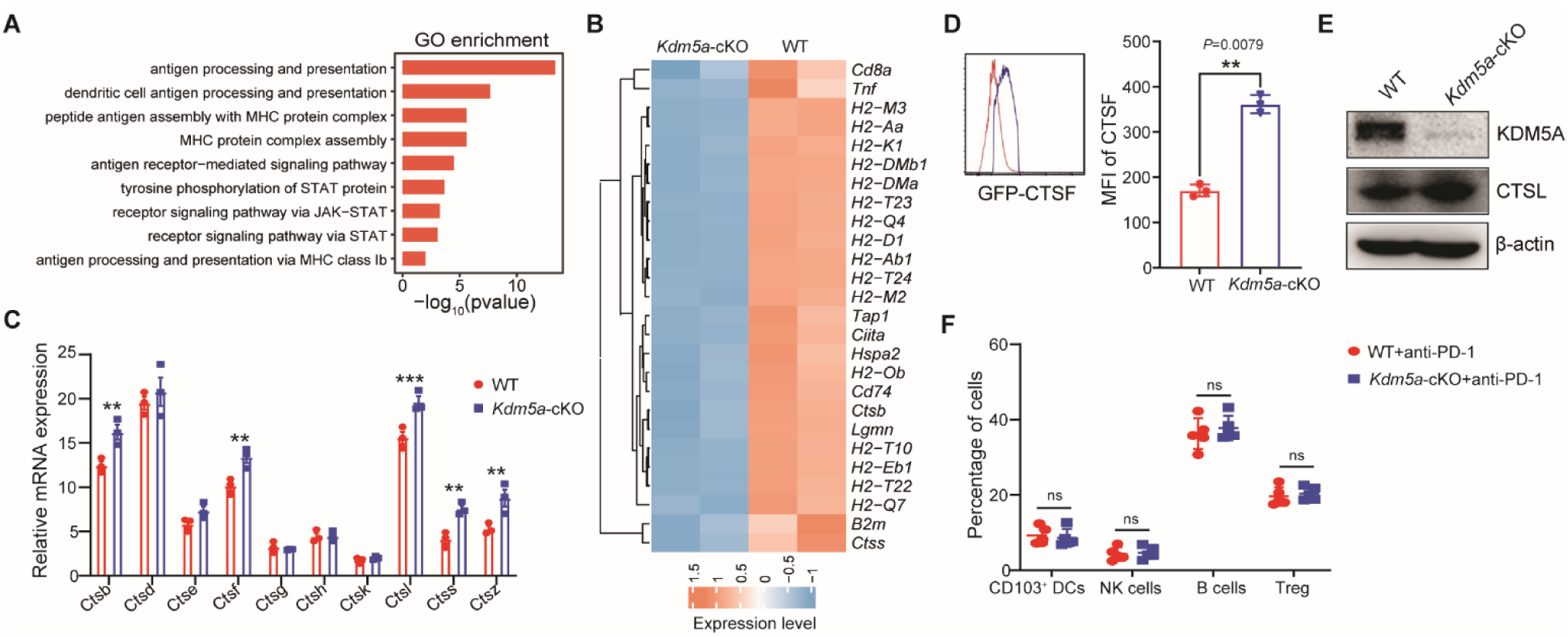
KDM5A directly targets and down-regulates lysosomal cathepsins in DCs. Related to Figure 5. **(A)** GO enrichment analysis based on the RNA-seq data of WT and *Kdm5a*-cKO BMDCs. **(B)** Heatmap of a hierarchical clustering of the mRNA level in the “antigen processing and presentation” cascade in WT and *Kdm5a*-cKO BMDCs based on RNA-seq data. Cluster analysis was performed with DAVID bioinformatics resources. **(C)** qPCR analysis of mRNA expression for relative lysosomal cysteine protease family genes in WT and *Kdm5a*-cKO BMDCs. The qPCR data were normalized to *β-actin*. **(D)** Representative flow cytometry analysis and quantification of CTSF in WT and *Kdm5a*-cKO cDC1. **(E)** Western blot analysis of CTSL in WT and *Kdm5a*-cKO cDC1 (B220^-^CD11c^+^MHCII^+^CD24^+^CD172a^-^) isolated from spleen. **(F)** Quantification by flow cytometry analysis of CD103^+^DCs, NK, B and Tregs (CD4^+^CD25^+^Foxp3^+^) in CD45^+^ TILs in MC38 tumors from mice receiving the indicated treatments. Data are represented as mean ± s.e.m. One of two or three representative experiments was shown (C, D-F). Statistical analysis was performed using two-sided unpaired Student’s *t*-test (C, D, F). \**P* < 0.05, \*\**P* < 0.01, and \*\*\**P* < 0.001.

**Figure S7.**
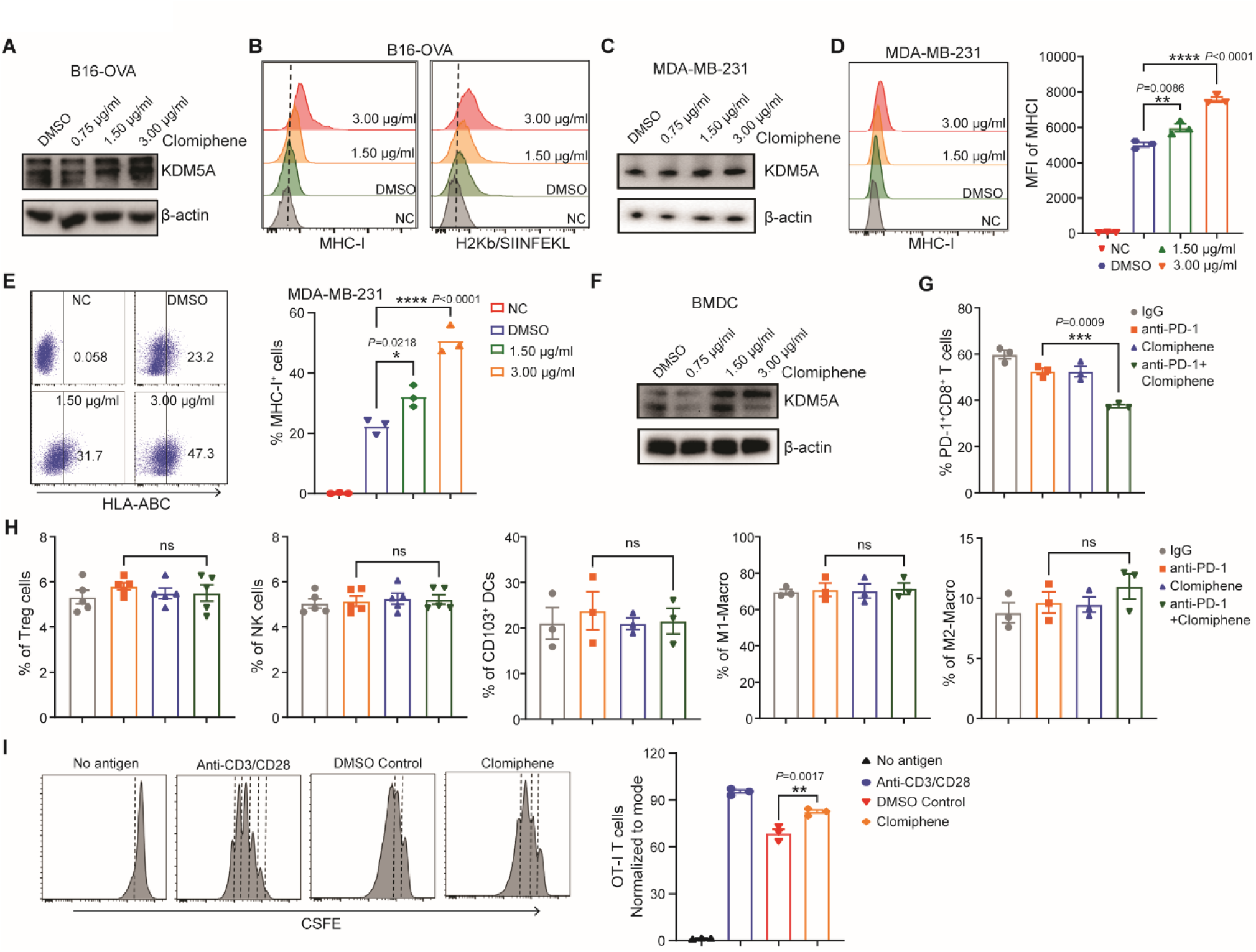
Clomiphene increases KDM5A expression in both tumor cells and DCs both *in vitro* and *in vivo*. Related to Figure 6. **(A)** Western blot analysis of KDM5A in B16-OVA cells treated with clomiphene (0.75, 1.5 and 3 μg/ml). **(B)** MHC I and H2Kb/SIINFEKL were visualized by flow cytometry on negative control (NC), DMSO, and Clomiphene (1.5 and 3 μg/ml) treated B16-OVA cells. **(C)** Western blot analysis of KDM5A in MDA-MB-231 cells treated with clomiphene (0.75, 1.5 and 3 μg/ml) **(D-E)** MHC-I and H2Kb/SIINFEKL were visualized by flow cytometry on negative control (NC), DMSO, and Clomiphene (1.5 and 3 μg/ml) treated MDA-MB-231 cells. **(F)** Western blot analysis of KDM5A in BMDCs treated with clomiphene (0.75, 1.5 and 3 μg/ml for 24 hr). **(G)** Quantification by flow cytometry analysis of PD-1^+^CD8^+^ T cells in B16-OVA tumors at day 18 of indicated treatments as Figure 6B. **(H)** Quantification by flow cytometry analysis of Tregs, NK cells, CD103^+^ DCs, M1-macro and M2-macro in B16-OVA tumors with the indicated treatments. **(I)** OT-I CD8^+^ T cell proliferation stimulated by no antigen (negative control), anti-CD3/anti-CD28 (positive control), DMSO control and clomiphene (3 μg/ml) treated B16-OVA cells. Data are represented as mean ± s.e.m. One of two or three representative experiments was shown. Statistical analysis was performed using one-way ANOVA with Bonferroni’s multiple comparison tests (D, E, G, H, I). \**P* < 0.05, \*\**P* < 0.01, \*\*\**P* < 0.001, and \*\*\*\**P* < 0.0001.

